# Transcriptional development of phospholipid and lipoprotein metabolism in different intestinal regions of Atlantic salmon (*Salmo salar*) fry

**DOI:** 10.1101/194514

**Authors:** Yang Jin, Rolf Erik Olsen, Mari-Ann Østensen, Gareth Benjamin Gillard, Sven Arild Korsvoll, Nina Santi, Arne Bjørke Gjuvsland, Jon Olav Vik, Jacob Seilø Torgersen, Simen Rød Sandve, Yngvar Olsen

## Abstract

**Background:** It has been suggested that the high phospholipid (PL) requirement in Atlantic salmon (*Salmo salar*) fry is due to insufficient intestinal *de-novo* synthesis causing low lipoprotein (LP) production and reduced transport capacity of dietary lipids. However, there has not been performed any in-depth ontological analysis of intestinal PL and LP synthesis with development of salmon. Therefore in this paper we used RNA-seq technology to test the hypothesis that the high PL requirement in salmon fry was associated with undeveloped PL synthesis and LP formation pathways in intestine. There was a special focus on the understanding homologous genes, especially from salmonid-specific fourth vertebrate whole-genome duplication (Ss4R), contribution to salmonid specific features of regulation of PL metabolic pathways. The study was performed in stomach, pyloric caeca and hindgut at 0.16g (1 day before first-feeding), 2.5g and 10g of salmon.

**Results:** In general, we found an up-regulation of *de-novo* phosphatidylcholine (PtdCho) synthesis, phosphatidylethanolamine (PtdEtn) and LP formation pathways in pyloric caeca of salmon between 0.16g and 10g. Thirteen genes in these pathways were highly (*q*<0.05) up-regulated in 2.5g salmon compared to 0.16g, while only five more significant (*q*<0.05) genes were found when the fish grew up to 10g. Different homologous genes were found dominating in stomach, pyloric caeca and hindgut. However, the expression of dominating genes in PL and LP synthesis pathways was much higher in pyloric caeca than stomach and hindgut. Salmon-specific homologous (Ss4R) genes had similar expression during development, while other homologs had more diverged expression.

**Conclusions:** An increasing capacity for PL synthesis and LP formation was confirmed in pyloric caeca. The up-regulation of the *de-novo* PtdCho pathway confirms that the salmon fry have increasing requirement for dietary PtdCho compared to adult. The similar expressions between Ss4R homologous genes suggest that the functional divergence of these genes was incomplete compared to homologs derived from other whole genome duplication. The results of the present study have provided new information on the molecular mechanisms of phospholipid synthesis and lipoprotein formation in fish.

## Background

Phospholipids (PL) are the main constituent of all biological cell membranes and separate the intracellular and extracellular aqueous environment. In addition to providing a structural scaffold for cell membranes, PL are also involved in numerous biological functions, such as provision of metabolic energy, cell membrane transport, and regulation of metabolism [1]. PL are also key structural components of lipoproteins (LP), which are involved in transport of dietary lipids from intestinal enterocytes to liver and peripheral tissues [2, 3]. It is well known that dietary inclusions of PL can improve growth and performance in many fish species including Atlantic salmon *Salmo salar* [4], Atlantic cod *Gadus morhua* [5] and rainbow trout *Oncorhynchus mykiss* [6]. Dietary and biliary PL is mainly digested by intestinal phospholipase A_2_ in fish resulting in 1-acyl lyso-phospholipids (lyso-PL) and free fatty acids (FFA). Subsequently, both lyso-PL and FFA are absorbed into intestinal enterocytes, re-esterified into PL before being exported to the rest of the body [7]. In addition, PL can be synthesized *de-novo* in enterocytes from glycerol-3-phosphate (G-3-P) in fish [8].

A minimum requirement of PL is associated with early developmental stages of fish, but so far no minimum requirement has been demonstrated in adult fish [8]. In line with this, dietary PL has been shown to enhance growth and survival in Atlantic salmon fry up to 2.5 g, but not in larger fish [4, 9]. It has been suggested that the higher PL requirement is due to insufficient ability of *de-novo* synthesis in the intestine leading to low LP production and consequently reduced transport capacity of dietary lipids [8]. This was supported by previous histological studies in salmonids, showing lipid accumulation in intestinal enterocytes when fed PL deficient diet [9, 10]. However, these differences were not evident in a previous transcriptomic study where gene expressions in PL biosyntheiss pathways were not changed in 2.5g salmon fed by PL-supplemented diet [11]. No study has demonstrated the gene expression in salmon smaller than 2.5g.

The intestinal tract of salmon consists of several regions, with different functions in lipid digestion, absorption and transport. It is generally believed that pyloric caeca (PC), rather than stomach (SM) or hindgut (HG), is the predominant region for lipid absorption and transport in salmon [12, 13]. Therefore, PL and LP were assumed to be mostly synthesized in PC region. However, other tissues like SM and HG could also have ability of synthesizing PL due to its structural roles in cell membranes [1]. LP has occasionally been observed in hindgut, suggesting some lipid absorption and transport activities in the region [14]. So far no study has demonstrated the PL metabolic pathways in SM and HG of fish.

Many homologous genes in mammals were found belong to gene families controlling the same enzymatic process but have distinct regulation in different tissues and developmental stages [1]. In this respect the Atlantic salmon has another layer of functional genome complexity as it experienced two extra rounds of whole genome duplication (WGD) compared to mammals, at the base of all teleost (Ts3R) and in a common ancestor of all salmonids ∼100-80 Mya (Ss4R) [15, 16]. Of the Ts3R and Ss4R gene duplicates, ∼20 and 55%, respectively, are still retained as expressed genes in the genome [17]. This dramatic increase in the number of homologous genes in salmon thus necessitates a careful annotation of PL synthesis and LP formation pathway genes and their tissue-specific expression regulation to improve our understanding of salmon PL metabolism.

In this paper we annotate and characterize gene regulation involved in PL synthesis and LP formation in different intestinal regions (SM, PC and HG) during early developmental stages of salmon. Our aims are to (I) improve our understanding of the homologous genes, especially from Ss4R, contribution to salmonid specific features of regulation of PL metabolism pathways and to (II) specifically test the hypotheses that PL requirements in early-developmental stages of Atlantic salmon is associated with insufficient PL synthesis and LP formation pathways.

## Methods

### Fish, diet and sampling procedure

Atlantic salmon eggs were hatched and cultivated at AquaGen Breeding Centre (Kyrksæterøra, Norway). From first-feeding, the fish were fed a normal commercial diet which satisfies the nutritional requirement of salmon, but without any additional PL supplement (Additional file 1). The diet was produced by EWOS AS (Bergen, Norway). Three salmon individuals (n=3) were sampled at sizes of 0.16g (1 day before first feeding), 2.5g (65 days after first feeding) and 10g (100 days after first feeding). The fish were euthanized by 1g/L MS-222 (FINQUEL, Argent chemical labs, Washington, USA) buffered with same amount of sodium bicarbonate before dissection. Samples of SM, PC and HG were dissected and immediately placed in 1mL RNALater. Tissues were stored for 24h at 4°C for sufficient penetration of RNALater before being transferred to -80°C for long-term storage.

### RNA extraction, library preparation and transcriptome sequencing

The RNA extraction and library preparation were carried out in Centre for Integrative Genetics (CIGENE, Ås, Norway). Total RNA was extracted from SM, PC and HG using RNeasy Plus Universal Kits (QIAGEN, Hilden, Germany), according to manufacturer’s instruction. RNA concentration and purity were assessed by Nanodrop 8000 (Thermo Scientific, Wilmington, USA). RNA integrity was checked by Agilent 2100 Bioanalyzer (Agilent Technologies, Santa Clara, CA, USA). All samples have RIN value >8, which were sufficient for transcriptome analysis. RNA libraries were prepared by using TruSeq Stranded mRNA Library Prep Kit (Illumina, San Diego, CA, USA), according to manufacturer’s instruction. Samples were sequenced using 100bp single-end high-throughput mRNA sequencing (RNA-seq) on Illumina Hiseq 2500 (Illumina, San Diego, CA, USA) in Norwegian Sequencing Centre (Oslo, Norway).

### Identification of genes and phylogenetic analysis

The genes involved in PL synthesis and LP formation in Atlantic salmon (*Salmo salar*) were manually annotated by matching salmon proteins to zebrafish (*Danio rerio*) orthologs from the KEGG reference pathway of glycerolphospholipid metabolism and other studies [18-20]. Ortholog group predictions were carried out using Orthofinder (v0.2.8) on proteins from seven fish species: zebrafish (*Danio rerio*), stickleback (*Gasterosteus aculeatus*), medaka (*Oryzias latipes*), Northern Pike (*Esox lucius*), grayling (*Thymallus thymallus*), rainbow trout (*Oncorhynchus mykiss*), Atlantic salmon (*Salmo salar*), and two mammal outgroup species: human (*Homo sapiens*) and mouse (*Mus musculus*). The protein sequences within orthogroups were aligned to each other using MAFTT and maximum likelihood trees were estimated using FastTree. Orthogroup trees were subsequently split into smaller clan trees using an in-house R script (clanfinder.R, available from github). For zebrafish proteins in selected KEGG reference pathways, salmon proteins within the same protein clan tree were annotated using the zebrafish KEGG Orthology terms. The detailed information on Orthogroup prediction and phylogenetic analysis was published elsewhere [19]. All annotated salmon genes was grouped into a PL gene list and used for gene expression analysis.

### RNA-sequencing data and statistical analysis

Read sequences were quality trimmed, removing any Illumina TruSeq adapter sequence and low quality bases (Phred score < 20) from read ends and length filtered (minimum length 40 bases) using cutadapt (v1.8.1), before being aligned to the salmon genome (ICSASG_v2) using STAR (v2.5.2a). Raw gene counts per sample were generated from read alignments using HTSeq-count (v0.6.1p1) and the NCBI salmon genome annotation (available for download at http://salmobase.org/Downloads/Salmo_salar-annotation.gff3). The uniquely mapped reads, aligned to exon regions, were counted for each gene in the annotation.

For each tissue type (SM, PC, HG), a differential expression analysis (DEA) was performed comparing 2.5g and 10g samples to the 0.16g samples. Genes were filtered prior to DEA testing by a minimum count level of at least 1 count per million (CPM) in two or more samples, to remove genes with too few counts for testing. From raw counts, DEA was conducted using R package edgeR (v3.8.6) with pairwise exact tests to produce gene fold changes and *p* values. Genes with a false discovery rate (FDR) adjusted *p* value (*q*) < 0.05 were considered to be differentially expressed between two test conditions. The PL gene list was applied to retrieve DEA results for genes involved in PL and LP synthesis.

For comparing expression levels between different genes and tissues, normalized counts in the form of transcripts per million (TPM) values were generated. Raw gene counts were first divided by their mRNA length in kilobases to normalize for transcript length, then divided by the total number of counts from each library to normalize for sequencing depth.

All RNA-seq data analysis was preformed using R (v3.2.4) and Bioconductor (v3.3). The pathway maps of PL and LP synthesis were produced using PathVisio. The heatmap was drawn using R with package pheatmap. All other figures were produced using SigmaPlot for Windows Version 13.0.

## Results

### Annotation of PL pathway orthologs in salmon

In order to get comprehensive knowledge on the expression of genes involved in PL and LP synthesis in salmon, we created a PL gene list of all salmon genes in the pathways based on their zebrafish paralogs identified in previous studies. A total of 62 zebrafish genes involved in PL *de-novo* synthesis, lyso-PL synthesis, PL turnover and LP synthesis pathways were used to identify PL metabolism genes in salmon. In total, 125 corresponding salmon genes were identified and named based on their phylogenetic relationship to human and zebrafish homologs. Due to the Ss4R WGD, 67% of the PL genes contain a salmon-specific duplicate in the genome. Homologous genes controlling same enzymatic reaction were grouped into a family for comparison of gene expression. A summary of identification and nomenclature of PL genes is shown in Additional file 2.

### Tissue specific regulation of PL metabolism in the gut

The raw sequencing data are publicly available on European Nucleotide Archive under accession number PRJEB21981. An average total of 22 million reads were collected from each library, out of which ∼85% were mapped to the salmon genome ICSASG_v2 (Additional file 3). From a total of 81574 genes currently annotated, 31411 genes passed a minimum level of read counts for use in DEA. DEA was carried out on SM, PC and HG separately to assess the extent of developmentally associated changes in PL metabolism by comparing 2.5g and 10g salmon to 0.16g. The different intestinal regions differed greatly in the numbers of differentially expressed genes (DEGs, *q*<0.05), with 10% DEGs in SM, and around 30% DEGs in PC and HG (Additional file 4).

Relative expression of genes involved in PL synthesis and LP formation show categorization into three distinct tissue related clusters associated to developmental differences (Fig. 1). The genes in cluster 2 are characterized by having highest expression in PC, while the remaining genes were either highest expressed in SM (Cluster 1) or HG (Cluster 3). Only a few genes describe a reduced expression following development of fish, whereas most genes show onset of expression, especially in cluster 2. The DEGs in SM, PC and HG of 2.5g and 10g compared to 0.16g salmon were also annotated in Figure 1. Similar to the genome-wide changes in expression, PC and HG were much more responsive (40% DEGs) compared to SM (less than 20% DEGs). Most DEGs were shared between PC and HG in Cluster 2, while DEGs in other clusters were much fewer. Moreover, the shared DEGs showed a much larger change in PC than in HG during development, resulting in increasing difference of expression between PC and HG as salmon growth.

**Figure 1.**
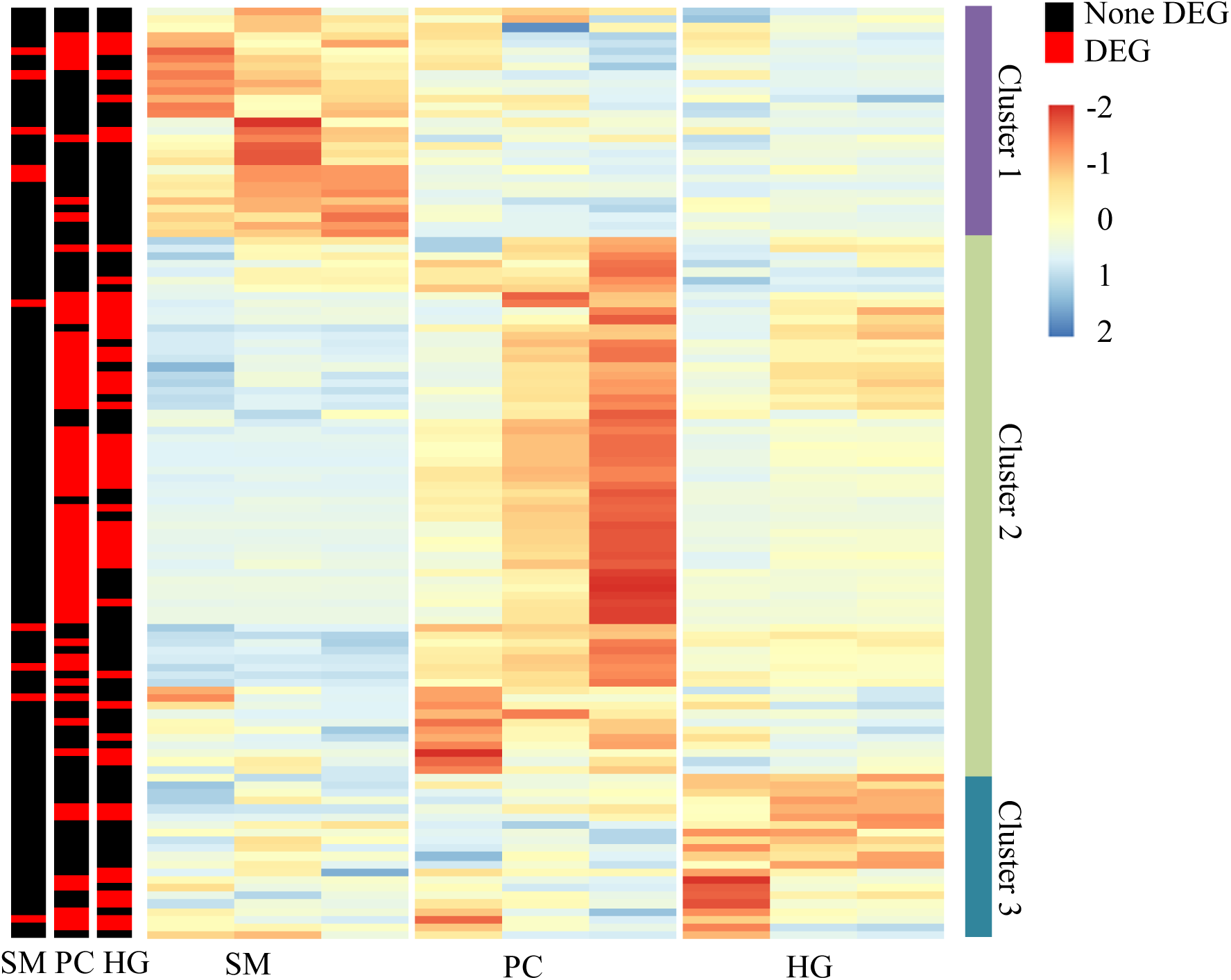
Expression of genes in phospholipid and lipoprotein synthsis pathways between different intestinal regions of salmon. For each tissue, the three columns represent 0.16g, 2.5g and 10g samples from left to right. The color intensity is relative to the standard deviation from mean of TPM over developmental stages and tissues (raw-scaled). Differential expressed genes (DEG, *q*<0.05) between 0.16g, 2.5g and 10g samples were annotated in three columns, which represent stomach (SM), pyloric caeca (PC) and hindgut (HG) respectively from left to right.

### Regulatory divergence of homologous genes among developmental stages and intestinal sections among Ss4R duplicates

The relative expression (TPM value) of all genes in the PL and LP synthesis pathways are summarized in Additional file 5. The salmon-specific (Ss4R) homologous genes showed mostly a similar divergence among developmental stages and intestinal sections. Moreover, the highest expression among the homologous genes was mostly found in PC rather than in SM or HG, which has supported that PC is the most important intestinal section for PL synthesis and LP formation in salmon.

*Pcyt1* family is a representative example to demonstrate the regulatory complexity of homologous genes among intestinal sections and developmental stages (Figure 2). The *pcyt1* family has 2 members in mammals (*pcyt1a* and *pcyt1b*), 4 members in zebrafish (*pcyt1aa*, *pcyt1ab*, *pcyt1ba* and *pcyt1bb*) and 7 members in salmon (*pcyt1aa*, *pcyt1ab_1*, *pcyt1ab_2*, *pcyt1ba_1*, *pcyt1ba_2*, *pcyt1bb_1* and *pcyt1bb_2*). Among all homologous genes, *pcyt1bb_1* had much higher expression levels in PC than in SM and HG. It was also the highest expressed homolog in all tissues. The expression level of *pcyt1bb_1* in PC and HG both more than doubled as the fish grew from 0.16g to 10g. The Ss4R homologous genes, *pcyt1bb_1* and *pcyt1bb_2*, showed similar divergence between tissues and developmental stages. The expressions of *pcyt1ab_1* and *pcyt1ba_1* genes were both higher in SM than PC and HG, while *pcyt1aa*, *pcyt1ab_2* and *pcyt1ba_2* were similarly expressed between the three tissues.

**Figure 2.**
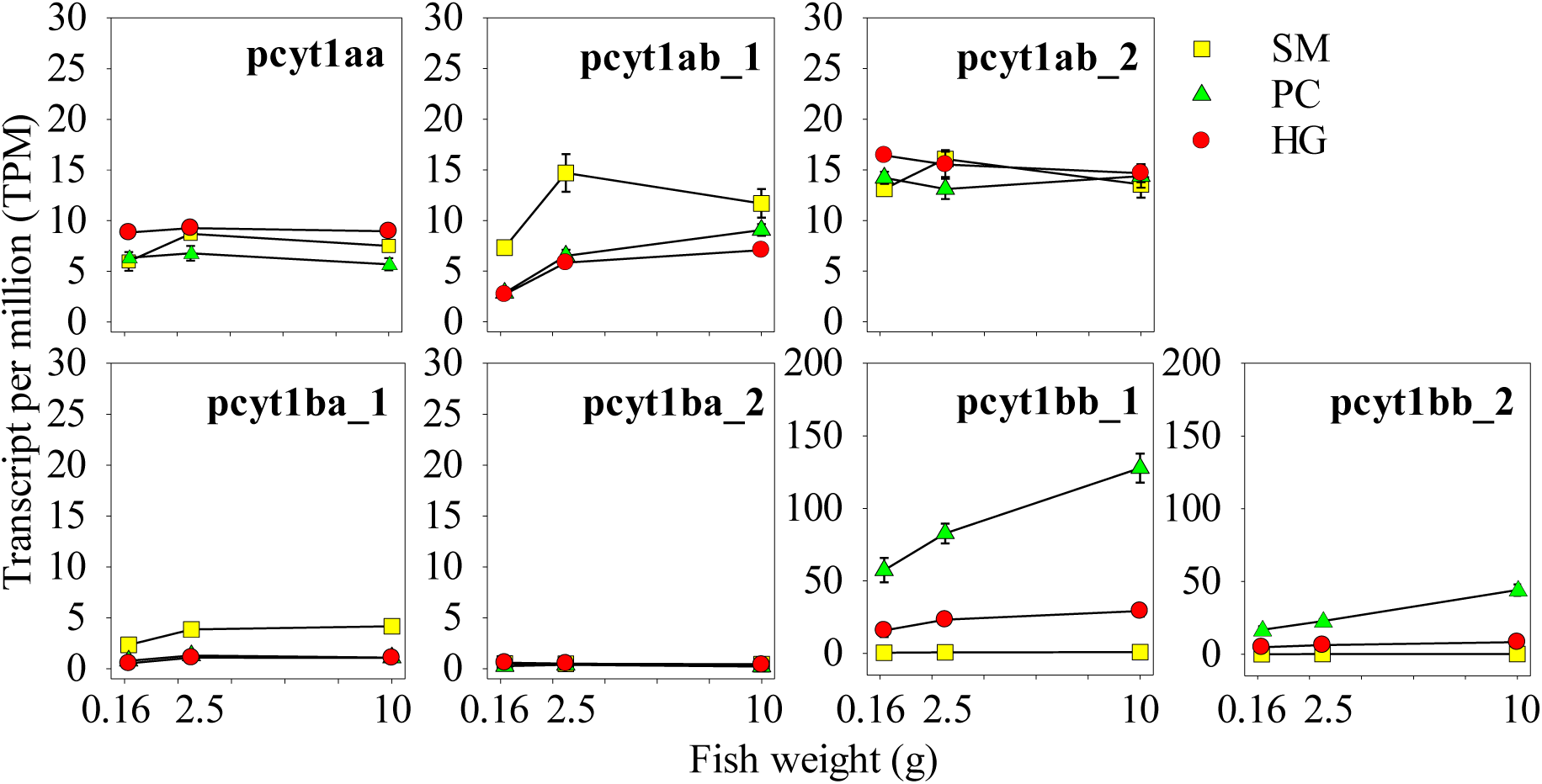
Comparison of *pcyt1* genes in TPM over developmental stages and intestinal regions. The gene expressions were compared in transcript per million (TPM) over three developmental stages (0.16g, 2.5g and 10g) in stomach (SM), pyloric caeca (PC) and hindgut (HG) of salmon. Numbers after underline indicates Ss4R gene duplicates specific in salmonids.

### Regulation of PL synthesis and LP formation pathways in PC

It clear from our expression analyses (Figure 1) that PC is the most active tissue, measured as expression level, with regards to PL and LP metabolic pathways. Therefore, PC was selected for a detailed study of the differences in expression regulation of homologous genes between 0.16g, 2.5g and 10g salmon (Figure 3). The homologous genes in eighteen families of key genes in PtdCho, PtdEtn and LP synthesis pathways were thus selected for in depth analyses.

**Figure 3.**
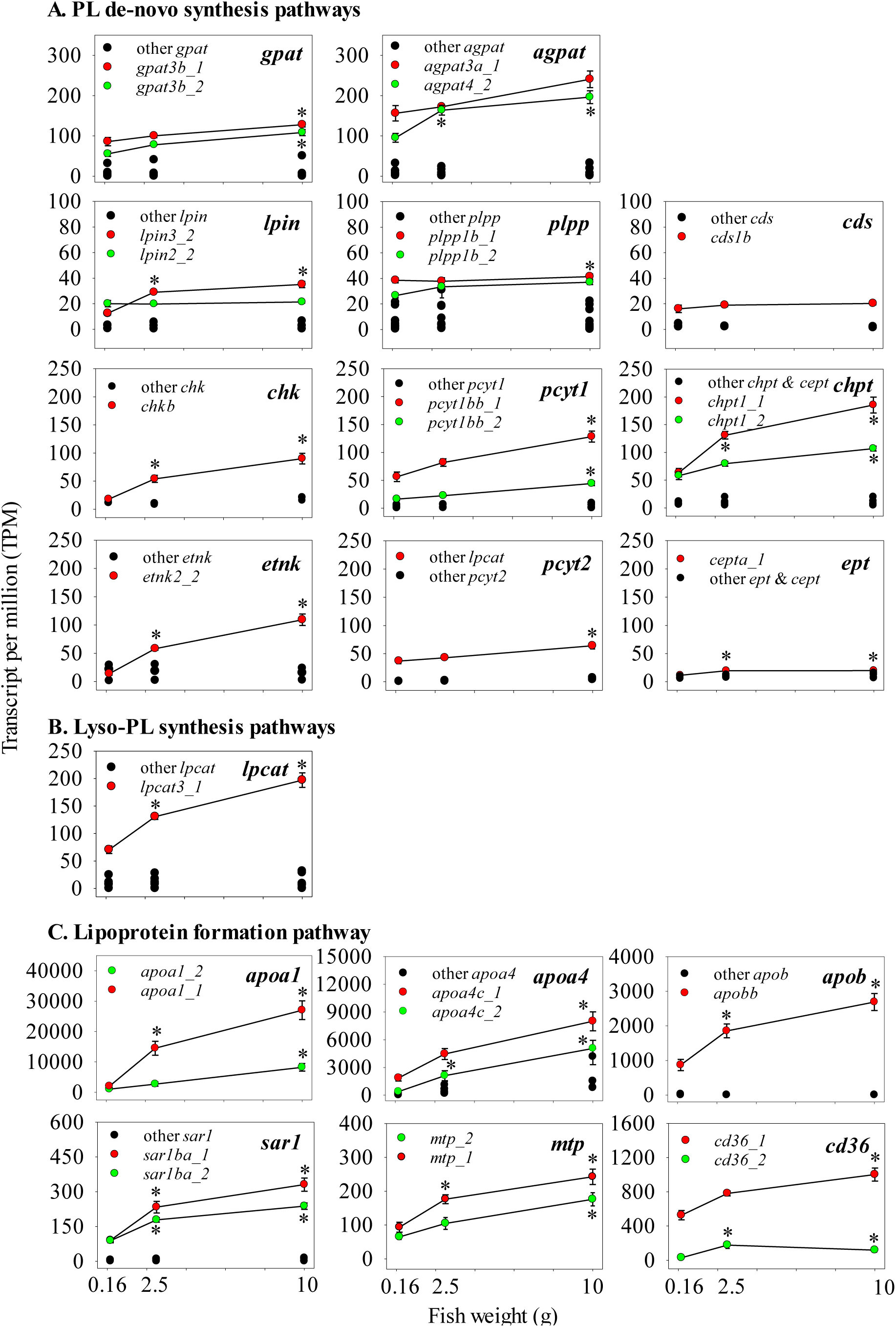
Expressions of key genes in phospholipid and lipoprotein synthesis pathways in pyloric caeca at early stage of salmon. Eighteen families of homologous genes in phospholipid (PL) *de-novo* synthesis, lyso-phospholipid (lyso-PL) synthesis and lipoprotein formation pathways are shown for comparing their expression between 0.16g, 2.5g and 10g fish. Genes with high TPM are marked in red and green, while other genes were all marked in black.

In most gene families, one or two genes had much higher expression than their homologs, with all being up-regulated during development (Figure 3). In the eleven families with two highly expressed genes, they are mostly salmon-specific duplicates from Ss4R WGD. In *de-novo* pathways, highly expressed genes were mostly significantly (*q*<0.05) up-regulated from 0.16g to 10g (Figure 3 A). The highly expressed genes involved in the PtdCho synthesis pathway (*chk*, *pcyt1* and *chpt* families) showed a much more pronounced increase compared to genes in other *de-novo* pathways. By comparing TPM of the highly expressed genes, *lpin* and *plpp* had slightly higher expression compared to *cds* family. Genes in *ept* family had much lower expression than other families in the pathway. In lyso-PL synthesis pathways, the expression of *lpcat3a* was largely increased (*q*<0.05) during development, whereas other *lpcat* genes remained stable (*q*>0.05, Figure 3 B). *Lpgat1a* and *lpiat1* in other families in lyso-PL pathways were also up-regulated (*q*<0.05) during development (Additional file 6). All other genes in lyso-PL synthesis pathways were expressed much lower than *lpcat3a*. Genes in LP formation pathway had much higher expression than in PL synthesis pathways (Figure 3 C). All highly expressed genes in LP formation pathway were largely up-regulated (*q*<0.05) between 0.16g and 10g salmon.

The proposed regulation of PL synthesis and LP formation pathways is summarized in Figure 4. By summarizing the change of the highly expressed genes in PC, we found up-regulations of *de-novo* PtdCho and PtdEtn synthesis pathways and down-regulations of *de-novo* PtdSer and PtdGro synthesis pathways between 0.16g and 10g salmon (Figure 4 A). Other *de-novo* synthesis pathways were not changed during development. The Lyso-cardiolipin synthesis pathway was not changed, while other lyso-PL synthesis pathways were all up-regulated. The phospholipid turnover pathways were not changed during development. The LP formation pathway was up-regulated between 0.16g and 10g (Figure 4 B).

**Figure 4.**
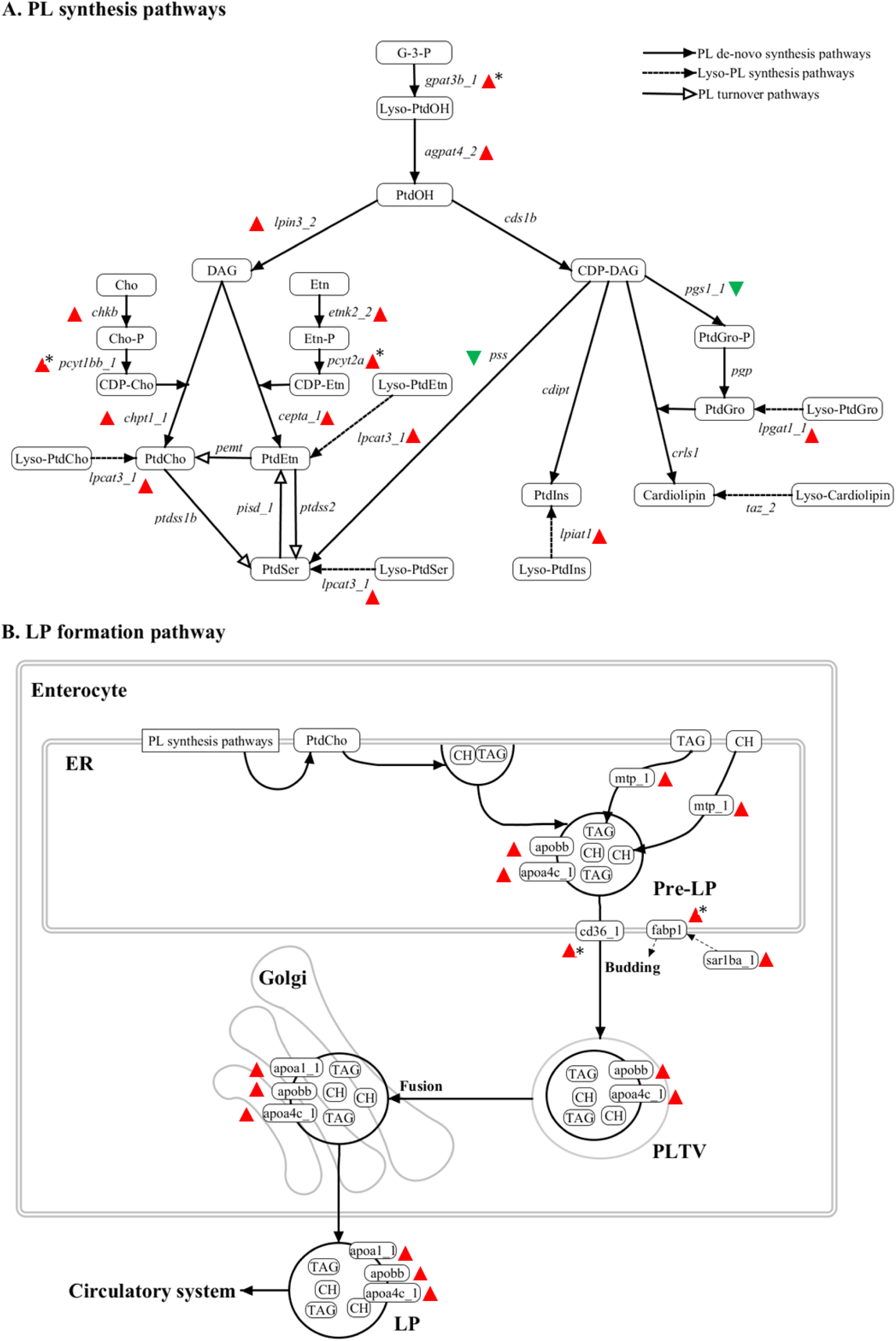
Comparison of phospholipid (PL) synthesis and lipoprotein (LP) formation pathways between 0.16g, 2.5g and 10g salmon. Colored triangles indicate the significantly (*q*<0.05) up (red) or down (green) regulation of the highest expressed genes found in each enzymatic step of the pathways. Asterix indicates genes only significantly (*q*<0.05) changed between 0.16g and 10g. A) PL *de-novo* synthesis, lyso-PL synthesis and PL turnover pathways in fish. Glycerol-3-phosphate (G-3-P) is first acylated by acyltransferases to phosphatidic acid (PtdOH), which can be transferred into diacylglycerol (DAG) or CDP-diacylglycerol (CDP-DAG) by phosphatidate phosphatase (plpp and lpin) or CDP-DAG synthetase (cds). DAG is utilized with CDP-choline (CDP-Cho) and CDP-ethanolamine (CDP-Etn) for synthesizing of phosphatidylcholine (PtdCho) and phosphatidylethanolamine (PtdEtn). CDP-DAG is utilized for synthesizing of phosphatidylserine (PtdSer), phosphatidylglycerol (PtdGro), phosphatidylinositol (PtdIns) and Cardiolipin. **B)** LP formation pathway in enterocyte of fish. PtdCho is synthesized on the membrane of endoplasmic reticulum (ER) through de-novo synthesis, turnover or lyso-PL pathway before used for pre-lipoprotein (Pre-LP) formation. Pre-LP is a nascent lipoprotein assembled by PtdCho, triacylglycerol (TAG), cholesterol (CH), apolipoprotein B (apob) and apolipoprotein AIV (apoa4). Pre-LP is then targeted to the Golgi apparatus via pre-lipoprotein transport vesicle (PLTV) generated by ER. The maturation of Pre-LP happens in Golgi, where apolipoprotein AI (apoa1) is added before secreting into circulatory system.

## Discussions

The objective of the present study was to explore the transcriptional changes of PL synthesis and LP formation pathways in different intestinal regions of salmon during early developmental stages. By integrating RNA-seq data with a manually curated automated sequence ortholog prediction, we identified many DEGs in SM, PC and HG during development of salmon. Most of those changes occurred between start feeding at 0.16g and 2.5g, and continued up to 10g. This shows that the maturation of lipid metabolic pathways progressed for a substantial period following the completion of yolk sac reabsorption. By comparing the highly expressed genes in each family, we found a continuous increased capacity of PtdCho, PtdEtn and LP synthesis in PC from 0.16g to 10g. This implies an increased capacity for PL and LP synthesis after onset of start feeding.

Our results are in line with previous qPCR-based single gene studies in salmon where several major PL biosynthetic genes were up-regulated in PtdCho and PtdEtn synthesis between 2.5g and 10g [18]. However, we did not observe a clear levelling-off of expression at 10g, suggesting that the completion of intestinal maturation might take longer time to accomplish. This supports the hypotheses of a higher dietary PL requirement in salmon fry compared to larger fish previously put forward [10]. The present study was the first to investigate expression differences between homologous genes in PL and LP synthesis pathways in salmon. Several homologous genes were identified controlling the same enzymatic step in PL and LP synthesis. In most cases, we found that one or two genes had much higher expression levels than their homologs for a given tissue, suggesting these genes to be key regulators in the pathways. However, the expression levels of the homologous genes appear to vary with both intestinal regions and developmental stages. This resulted in different genes dominating in SM, PC and HG. The diverged expressions are likely due to different functions of homologous genes in the tissues, subcellular location and developmental stages [1]. More interestingly, the salmon-specific homologous genes derived from Ss4R WGD expressed similarly in different tissues and developmental stages, while other homologous genes seemed to have more diverged function. This may suggest that the functional divergence of Ss4R homologous genes was incomplete compared to homologs from other WGDs. This supports the recent study in which 55% of the Ss4R homologous genes were found to have similar expression levels among 15 tissues in salmon [17].

The present study was the first to utilize RNAseq to investigate roles of different intestinal regions in PL metabolism. PC is the predominant region for lipid absorption and transport in salmon [12, 13]. This is consistent with higher expression of genes involved in PtdCho, PtdEtn and LP synthesis pathways in PC rather than SM and HG. As PtdCho is the predominant lipid class forming the membrane fraction of LP, the higher expression of PtdCho synthesis pathway in PC verifies a high rate of LP production [3, 8]. This has been confirmed by histological observation of large lipid droplets accumulating in midgut enterocytes of PL-deficient fish, while little droplets were found in fish fed dietary PtdCho [9, 10]. The expression of genes involved in PL synthesis in SM and HG were most likely related to other biological function such as cell maintenance and metabolism. Almost no expression of LP formation genes were found in SM, agrees with the general observation that SM is not involved in lipid digestion, uptake and transport. However, despite being expressed at low levels, many genes of the LP formation pathways were found in HG, suggesting some capacity to absorb and transport lipid in this intestinal region [14].

Several highly expressed genes in PL and LP synthesis pathways of salmon do not conform to the known function and regulation of these pathways in mammals. For example, it is believed that choline kinase (CK) α enzyme is critical for PtdCho maintenance in most tissues in mammals, whereas CKß enzyme is only essential for muscle tissue [21, 22]. In salmon, the expression of the *chkb* gene, encoding for CKß, was significantly elevated in PC after start feeding, whereas *chka* genes were unchanged. Similarly, the expressions of *pcyt1a* genes were relatively low in PC during early development, while *pcyt1bb_1* notably increased in PC after start feeding. Therefore, we assume that the production of PtdCho for LP synthesis is probably through a compensatory pathway controlled by *chkb* and *pcty1b* genes and activated in PC after switching to external feeding. On the other hand, there may be another pathway controlled by *chka* and *pcyt1a* genes, which produce PtdCho to maintain cell growth and survival. This suggestion agrees with previous studies pointing to the subcellular location of the enzymes [23, 24]. The CTP: phosphocholine cytidylyltransferase (CCT) α, which is the product of the *pcyt1a* gene, is predominantly located in the nucleus. On the other hand, *pcyt1b*-encoded CCTßis localized in the endoplasmic reticulum (ER) and the cytosol, which could be utilized in synthesizing PtdCho for LP formation. However, as the level of gene expression does not always directly reflect relative importance of two similar enzymes in a pathway, the posttranslational modification like phosphorylation of CCT could also be critical in regulating the activity of enzymes without affecting the mRNA level [1].

## Conclusions

The present study has provided new information on the molecular mechanisms of PL synthesis and LP formation in salmon fry. By comparing the expression levels of homologous genes, we identified several genes which had highly expression among their homologs in PtdCho, PtdEtn and LP synthetic pathways in PC of salmon. Those highly expressed genes were all up-regulated during development, confirming the increasing capacity for PL synthesis and LP formation. Taken the lowered activity of PL genes in early life of salmon and the link to PL requirement, it seems likely that the highest requirement is during early first feeding, then reducing as the fish grows. The expression levels of the homologous genes in PL synthesis and LP formation pathways appeared to vary with both intestinal regions and developmental stages. This resulted in different genes dominating in SM, PC and HG. The salmon-specific homologous genes derived from Ss4R WGD expressed similarly among different tissues and developmental stages, while homologous genes from other WGD seemed to have more diverged function. More studies on both gene and protein levels are required to confirm these relationships during early stages of salmon development. Considering the present results on identification of key regulating genes in PL synthesis and lipid transport, we suggest a future study on the dietary requirement of PL at first-feeding stage, which focuses on the changes of key regulating genes involved in PL and LP synthesis pathways in PC of salmon.

## List of abbreviations

**agpat**, 1-acylglycerol-3-phosphate acyltransferases; **cd36**, cluster of differentiation 36; **CDP**, cytidine diphosphate; **cdipt**, cdp-diacylglycerol-inositol 3-phosphatidyltransferase; **CDP-DAG**, CDP-diacylglycerol; **CDP-Cho**, CDP-choline; **CDP-Etn**, CDP-ethanolamine; **cds**, CDP-DAG synthase; **cept**, CDP-choline/ethanolamine: diacylglycerol phosphotransferase; **CH**, cholesterol; **chk**, choline kinase; **Cho-P**, phosphocholine; **chpt**, CDP-choline: diacylglycerol phosphotransferase; **crls**, cardiolipin synthase; **etnk**, ethanolamine kinase; **ER**, endoplasmic reticulum; **ept**, CDP-ethanolamine: diacylglycerol phosphotransferase; **Etn-P**, phosphoethanolamine; **fabpl**, fatty acid binding protein, liver; **DAG**, diacylglycerol; **G-3-P**, glycerol-3-phosphate; **gpat**, glycerol-3-phosphate acyltransferase; **lclat**, lysocardiolipin acyltransferas**e; lpcat**, lysophosphatidylcholine acyltransferase; **mboat**, membrane bound o-acyltransferase; **mtp**, microsomal triglyceride transfer protein; **pcyt1**, choline-phosphate cytidylyltransferase; **pcyt2**, ethanolamine-phosphate cytidylyltransferase; **pemt**, phosphatidylethanolamine methyltransferase; **pgp**, phosphatidylglycero phosphatase; **pgs**, CDP-diacylglycerol: glycerol-3-phosphate phosphatidyltransferase; **pmt**, phosphoethanolamine n-methyltransferase; **plpp**, phosphatidate phosphatase; **pss**, CDP-diacylglycerol:serine phosphatidyltransferase; **ptdss**, phosphatidylserine synthase; **pisd**, phosphatidylserine decarboxylase; **PtdCho**, phosphatidylcholine; **PtdEtn**, phosphatidylethanolamine; **PtdGro**, phosphatidylglycerol; **PtdIns**, phosphatidylinositol; **PtdOH**, phosphatidic acid; **PtdSer**, phosphatidylserine; **sar1**, secretion associated, RAS related GTPase 1; **TAG**, triacylglycerol; **taz**, tafazzin.

## Declarations

### Ethics approval and consent to participate

The study was carried out within the Norwegian animal welfare act guidelines, in accordance with EU regulation (EC Directive 86/609/EEC) and approved by the Norwegian Animal Research Authority (NARA).

### Consent for publication

Not applicable

### Availability of data and materials

The raw gene counts data of the current study is available for download at <u>http://salmobase.org/Downloads/Salmo_salar-annotation.gff3</u>

The orthogroup prediction and phylogenetic analysis of salmon genes are included in this published article and its supplementary information files:

Gillard G, Harvey TN, Gjuvsland A, Jin Y, Thomassen M, Lien S, Leaver M, Torgersen JS, Hvidsten TR, Vik JO et al: Diet And Life Stage Associated Remodeling Of Lipid Metabolism Regulation In The Duplicated Atlantic Salmon Genome. bioRxiv 2017. <u>https://doi.org/10.1101/140442</u>

## Authors’ contributions

YJ, RO, NS and YO comprehended the idea and designed the experiments. YJ sampled fish, prepared RNA-seq and analyzed RNA-seq data, and was a major contributor in writing the manuscript. MØ assisted in fish sampling. SK assisted in fish rearing and maintenance. GG and SS assisted in RNA-seq data analysis. Annotation of salmon genes including orthogroup prediction and phylogenetic analysis were done by SS, AG, JV and JT. SS, RO and YO provided directions and critiques. All authors read and approved the final manuscript.

## Acknowledgements

Thanks to Dr. Maren Mommens and AquaGen AS for providing experimental facilities and other practical information. Thanks to Dr. Hanne Hellerud Hansen and Cigene for RNAseq analysis.

## Additional files

**Additional file 1: Table 1** Composition and nutritional value of the diet used in current experiment. (DOCX 15.5 kb)

**Additional file 2: Table 1** List of Atlantic salmon (Ssa) genes involved in phospholipid (PL) *de-novo* synthesis, lyso-PL synthesis and lipoprotein (LP) formation pathways. Nomenclature of salmon genes was based on their human (Hsa) and zebrafish (Dre) paralogs. Numbers after underline in Ssa names indicate salmon-specific gene duplicates. NCBI gene ID of salmon and zebrafish is also listed in table. Reference listed the origin of zebrafish genes used for identification of salmon genes. (DOCX 31.5 kb)

**Additional file 3: Table 1** Summary of mapping statistics of all 45 samples used for RNA sequencing (3 replicates * 3 tissue in 0.16g fish, 6 replicates * 3 tissue in 2.5g and 10g fish). (DOCX 16 kb) **Additional file 4: Table 1** Number of significantly (q<0.05) differentially expressed genes (DEG) between 2.5g, 10g and 0.16g salmon. (DOCX 16.1 kb)

**Additional file 5: Table 1** Transcript per million (TPM) of all gene duplicates in phospholipid and lipoprotein synthesis pathways in stomach, pyloric caeca and Hindgut of 0.16g, 2.5g and 10g salmon. (DOCX 32.8 kb)

**Additional file 6: Table 1** Log2 fold change (LogFC) and adjusted *p* value (*q*) of all gene duplicates in phospholipid and lipoprotein synthesis pathways in stomach, pyloric caeca and hindgut of 2.5g and 10g salmon both compared to 0.16g. (DOCX 35 kb)

